# Cortical tracking of speech in noise accounts for reading strategies in children

**DOI:** 10.1101/2020.01.16.907667

**Authors:** Florian Destoky, Julie Bertels, Maxime Niesen, Vincent Wens, Marc Vander Ghinst, Jacqueline Leybaert, Marie Lallier, Robin A. A. Ince, Joachim Gross, Xavier De Tiège, Mathieu Bourguignon

## Abstract

Humans’ propensity to acquire literacy relates to several factors, among which, the ability to understand speech in noise (SiN). Still, the nature of the relation between reading and SiN perception abilities remains poorly understood. Here, we dissect the interplay between (i) reading abilities, (ii) classical behavioral predictors of reading (phonological awareness, phonological memory and lexical access), and (iii) electrophysiological markers of SiN perception in 99 elementary school children (26 with dyslexia). We demonstrate that cortical representation of phrasal content of SiN relates to the development of the lexical (but not sublexical) reading strategy. In contrast, classical behavioral predictors of reading abilities and the ability to benefit from visual speech to represent the syllabic content of SiN account for global reading performance (i.e., speed and accuracy of lexical and sublexical reading). Finally, we found that individuals with dyslexia properly integrate visual speech information to optimize processing of syntactic information, but not to sustain acoustic/phonemic processing. These results clarify the nature of the relation between SiN perception and reading abilities in typical and dyslexic child readers, and identified novel electrophysiological markers of emergent literacy.

## Introduction

Acquiring literacy is tremendously important in our societies. Central for reading acquisition are adequate phonological awareness,^1–3^ phonological memory,^4,5^ and lexical access.^6–8^ The adequacy of the learning environment also plays a major role.^9,10^ In particular, the presence of recurrent noise in the learning environment can substantially hinder reading acquisition.^11,12^ Therefore, the ability to understand speech in noise (SiN)—which is known to differ between individuals ^13,14^—should modulate the negative impact of environmental noise on reading acquisition. And indeed, the quality of brainstem responses to syllables in noise predicts reading abilities and its precursors.^15^ Moreover, individuals with dyslexia often exhibit a SiN perception deficit,^16,17^ that is particularly apparent when the background noise is composed of speech.^18^ This deficit has been hypothesized to root in a deficit in phonological awareness,^19,20^ but contradictory reports do exist.^21^ The question of whether SiN processing abilities relate to reading due to a common dependence on classical behavioral predictors (i.e., phonological awareness, phonological memory and lexical access), or due to other cognitive or neurophysiological processes specific to SiN processing, is thus open. Furthermore, it is also unexplored which aspects of reading and SiN processing abilities are related. Understanding these relations is especially important given that acoustic noise is ubiquitous, and given how adverse dyslexia can be for the cognitive and social development of children.

Reading is a multifaceted process. Hence, it is reasonable to think that SiN processing might relate to a restricted set of aspects of reading. Following the Dual Route Cascaded (DRC) model, reading is supported by two different routes: the *sublexical* and the *lexical* routes.^22,23^ The sublexical route implements the grapheme to phoneme conversion. It is used when reading unfamiliar words or pseudowords, but it is not suitable to correctly read irregular words (*i.e., yacht*) and to acquire fluent reading. Skilled reading relies on the lexical route that supports fast recognition of the orthographic word form of familiar words. The lexical route is indispensable to read irregular words, benefits reading of regular words, and does not contribute to reading pseudowords. Remarkably, the brain would implement these two reading strategies in two distinct neural pathways.^24–27^

There are also several distinct aspects of SiN processing that could relate to reading, and these can be derived from electrophysiological recordings of brain activity during connected speech listening. When listening to connected speech, human auditory cortical activity tracks the fluctuations of speech temporal envelope at frequencies matching the speech hierarchical linguistic structures, i.e., phrases/sentences (0.2–1.5 Hz) and words/syllables (2–8 Hz).^28–38^ Such cortical tracking of speech (CTS) is thought to be essential for speech comprehension.^31,33,35,37,39–41^ Corresponding brain oscillations would subserve the segmentation or parsing of incoming connected speech to promote speech recognition.^31,32,37,39,42^ In SiN conditions, children and adults’ brain preferentially tracks the attended speech rather than the global auditory scene, though with reduced fidelity when the noise hinders comprehension.^28,29,38,43–53^ Assessing CTS in noise can therefore provide objective measures of the impact of noise on the cortical representation of the different hierarchical linguistic structures of speech. Also relevant is how SiN perception is impacted by noise properties. In essence, the relevant parameters for an acoustic noise in SiN conditions are the degree of energetic and informational masking.^54^ The noise is energetic when its spectrum is similar to that of speech, and non-energetic otherwise. The noise is informational when it is made up of other speech signals, and non-informational otherwise. An energetic noise introduces *physical* interferences and an informational noise introduces *perceptual* interferences. Finally, to enhance SiN processing, humans also benefit from visual information of the speaker’s articulatory mouth movements.^55,56^ All these aspects of SiN perception can be captured by measures of CTS.

In this study, we investigated the relations between reading abilities, neural representations of SiN quantified with CTS, and classical behavioral predictors of reading in elementary school children. To fully characterize cortical SiN processing, we measured CTS in several types of background noises introducing different levels of energetic and informational masking and in conditions where the face of the speaker was visible (*lips*) or not (*pics*) while talking. This study was designed to answer three major questions: (i) What aspects of cortical SiN processing and reading abilities are related in typically-developing elementary school children (ii) To what extent are these relations mediated by classical behavioral predictors of reading? (iii) Do these relations translate to alterations in dyslexic children in comparison with typical readers matched for age or reading-level? As preliminary steps to tackle these questions, we identify relevant features of CTS in noise, and assessed in a global analysis the nature of the information about reading brought by all the identified features of CTS in noise and classical behavioral predictors of reading abilities.

## Results

We first report on 73 children with typical reading abilities. Then, we report on 26 children with dyslexia matched with a sub-sample of the 73 typical readers for age (n = 26) or reading level (n = 26). Both control groups were included to tell whether development or reading experience can explain potentially uncovered SiN deficits.^57^ Reading performance and its classical behavioral predictors were characterized in a comprehensive cognitive evaluation (Table 1). Children’ brain activity was recorded with magnetoencephalography (MEG) while they were attending to 4 videos of ~6 min each (Figure 1). Each video featured 9 conditions: 1 noiseless and 8 SiN resulting from the combination of energetic or non-energetic and informational or non-informational noise with *lips* and *pics* visual inputs. For each condition, we regressed the temporal envelope of the attended speech on MEG signals with several time lags using ridge regression and cross-validation (see methods section for details).^58^ The ensuing regression model was used to reconstruct speech temporal envelope from the recorded MEG signal. CTS was computed as the correlation between the genuine and reconstructed speech temporal envelopes. We did this for MEG and speech envelope signals filtered at 0.2–1.5 Hz (phrasal rate)^28,59^ and 2–8 Hz (syllabic rate),^47,51,60,61^ and for MEG sensor signals in the left and right hemispheres separately.

**Figure 1.**
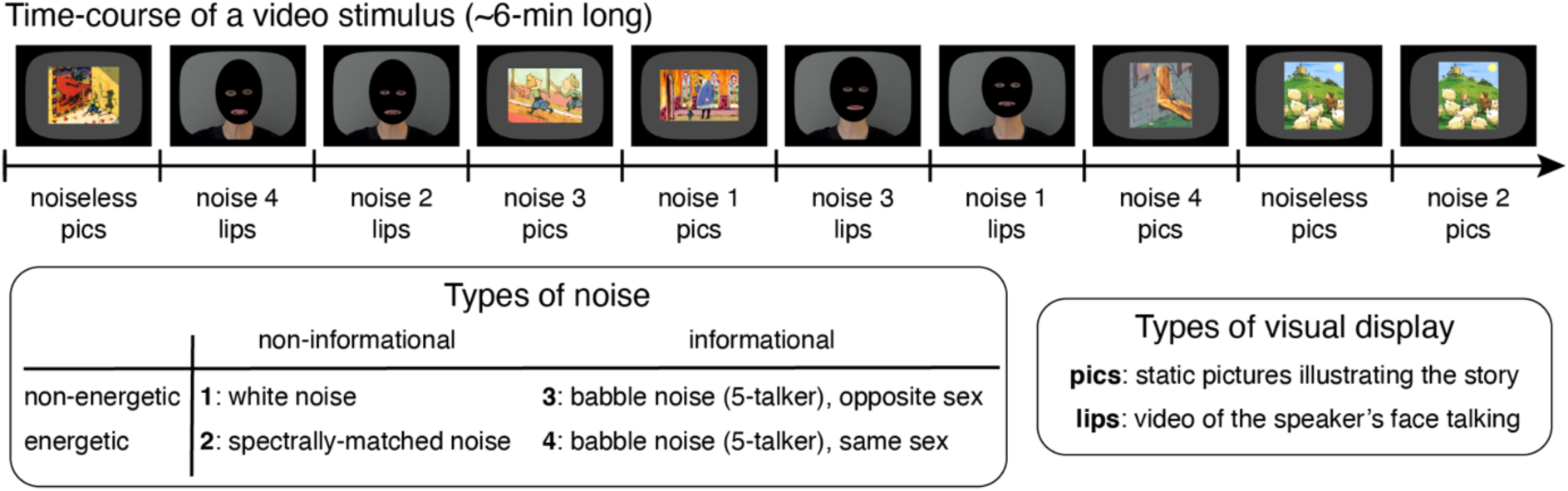
Illustration of the experimental material used in the neuroimaging assessment. Subjects viewed 4 videos of ~6 min duration. Each video was divided into 10 blocks to which experimental conditions were assigned. There were 2 blocks of the *noiseless* condition, and 8 blocks of speech-in-noise (SiN) conditions: 1 block for each possible combination of the 4 types of noise and two types of visual display. The interference introduced by the noise was either energetic or not and informational or not. The visual display provided visual speech information (*lips*) or not (*pics*).

**Table 1.** Mean and standard deviation of behavioral scores in each reading group of 26 children and comparisons (*t*-tests) between groups. The number of degrees of freedom (*df*) was 50 for all comparisons except those involving auditory attention (TAP) scores (dyslexic readers vs. controls in age, *df* = 49; dyslexic readers vs. controls in reading level, *df* = 38) and socio-economic status (dyslexic readers vs. controls in age, *df* = 49; dyslexic readers vs. controls in reading level, *df* = 47). IQ, intelligence quotient; RAN, rapid automatized naming; SD, standard deviation.

|  | Dyslexic readers |  | Age-matched control |  | Reading level control |  | Dyslexic readers compared with controls |  |  |  |
| --- | --- | --- | --- | --- | --- | --- | --- | --- | --- | --- |
|  | Mean | SD | Mean | SD | Mean | SD | in age |  | in reading level |  |
|  |  |  |  |  |  |  | p | t(*df*) | p | t(*df*) |
| Chronological age | 9.81 | 1.26 | 9.52 | 0.96 | 7.21 | 0.53 | 0.38 | 0.89 | **<0.0001** | **10.31** |
| Non-verbal IQ | 111 | 11 | 114 | 10 | 112 | 9 | 0.30 | -1.04 | 0.784 | -0.28 |
| Socio-economic status | 6.12 | 2.44 | 6.96 | 1.45 | 6.96 | 2.47 | 0.14 | -1.50 | 0.17 | -1.40 |
| Alouette reading accuracy | 89.0 | 5.7 | 96.2 | 2.1 | 89.0 | 6.46 | **<0.0001** | **-6.07** | 0.988 | 0.01 |
| Alouette reading speed | 141 | 61 | 292 | 91 | 138 | 64 | **<0.0001** | **-7.04** | 0.867 | 0.17 |
| Irregular words reading [words/s] | 0.54 | 0.33 | 1.16 | 0.44 | 0.40 | 0.35 | **<0.0001** | **-5.82** | 0.15 | 1.47 |
| Regular words reading [words/s] | 0.73 | 0.41 | 1.35 | 0.41 | 0.61 | 0.35 | **<0.0001** | **-5.51** | 0.29 | 1.06 |
| Pseudo-words reading [words/s] | 0.42 | 0.24 | 0.78 | 0.30 | 0.39 | 0.21 | **<0.0001** | **-4.88** | 0.61 | 0.50 |
| Visual attention | 30.3 | 3.74 | 32.0 | 2.69 | 27.4 | 4.43 | 0.070 | -0.95 | **0.014** | **2.53** |
| Phoneme suppression | 7.92 | 2.15 | 9.04 | 1.75 | 8.42 | 1.27 | **0.046** | **-2.05** | 0.313 | -1.02 |
| Phoneme fusion | 7.73 | 1.59 | 9.31 | 0.97 | 8.92 | 1.16 | **<0.0001** | **-4.32** | **0.003** | **-3.09** |
| Forward digit span | 5.08 | 0.84 | 5.8 | 0.69 | 5.15 | 0.78 | **0.001** | **-3.41** | 0.735 | -0.34 |
| Backward digit span | 3.69 | 0.79 | 4.5 | 1.33 | 3.38 | 0.75 | **0.011** | **-2.66** | 0.156 | 1.44 |
| RAN time [s] | 24.4 | 7.84 | 20.1 | 3.02 | 30.6 | 7.51 | **0.013** | **2.59** | **0.005** | **-2.91** |
| TAP mean response time [ms] | 627 | 99.0 | 613 | 75.4 | 667 | 93.4 | 0.59 | 0.53 | **0.07** | **-1.86** |
| TAP SD response time [ms] | 140 | 45.0 | 129 | 30.3 | 171 | 46.7 | 0.33 | 0.98 | **0.02** | **-2.36** |
| TAP correct responses | 15.6 | 0.58 | 15.7 | 0.68 | 15.3 | 1.07 | 0.42 | -0.81 | 0.11 | 1.65 |
| TAP false responses | 2.15 | 2.26 | 0.84 | 1.28 | 1.21 | 0.97 | **0.014** | **2.54** | 0.89 | 0.13 |

Table 2 presents the percentage of the 73 typical readers showing statistically significant phrasal and syllabic CTS, for both hemispheres, and each condition. All typical readers showed significant phrasal CTS in noiseless and non-informational (non-speech hereafter) noise conditions, and still most of them in informational (babble hereafter) noise conditions (mean ± SD across conditions, 98.3 ± 2.1 %). Most of the typical readers showed significant syllabic CTS in noiseless and non-speech noise conditions (93.8 ± 3.2 %), and slightly less of them in babble noise conditions (80.1 ± 4.3 %). This result clearly indicates that CTS can be robustly assessed at the subject level.

**Table 2.**
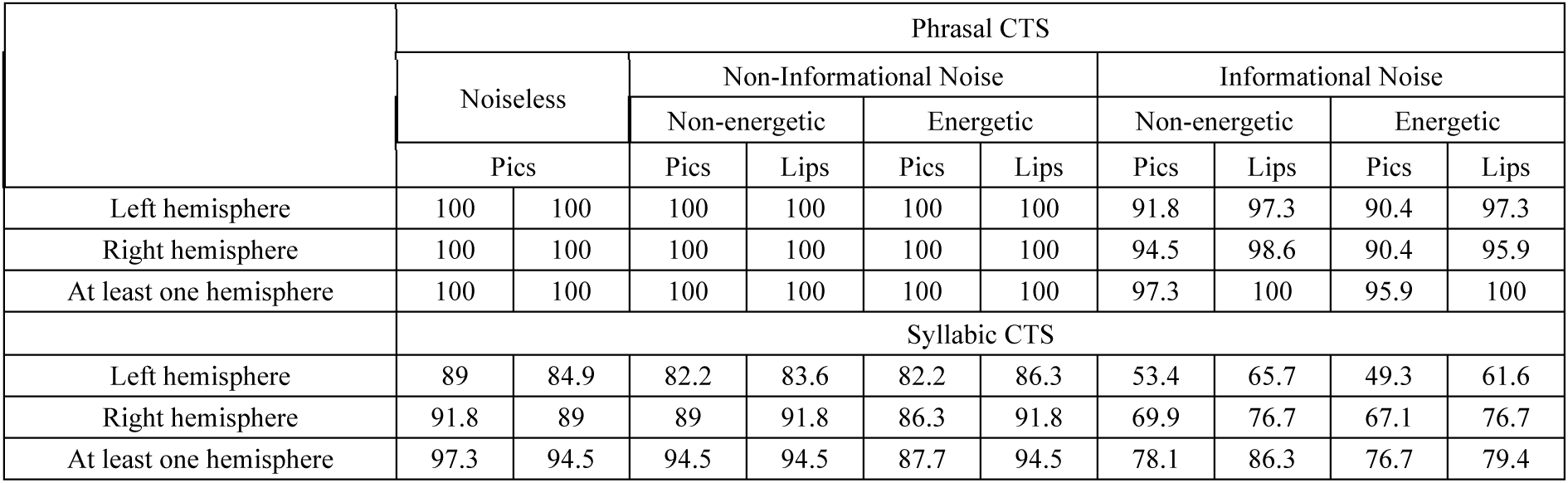
Percentage of the 73 typical readers showing significant cortical tracking of speech (CTS) at phrasal and syllabic rates in the 9 different conditions, in the left hemisphere (LH), in the right hemisphere (RH), or in at least one hemisphere. The two values provided for the noiseless condition correspond to two arbitrary subdivisions of the *noiseless* data to match the amount of data for the eight noise conditions.

### What aspects of SiN processing modulate the measures of CTS in noise?

First, we identify the main factors modulating CTS in SiN conditions. To that aim, we evaluated with linear mixed-effects modeling how the normalized CTS (nCTS) in SiN conditions depends on hemisphere, noise properties, and visibility of the talker’s lips. The nCTS is a contrast between CTS in SiN and noiseless conditions (see Methods) that takes values between –1 and 1, with negative values indicating that the noise reduces CTS. Such contrast presents the advantage of being specific to SiN processing abilities by factoring out the global level of CTS in the noiseless condition. In that analysis, nCTS values were corrected for age, time spent at school and IQ.

Table 3 presents the final linear mixed-effects model of phrasal and syllabic nCTS, and Figure 2 illustrates underlying values.

**Figure 2.**
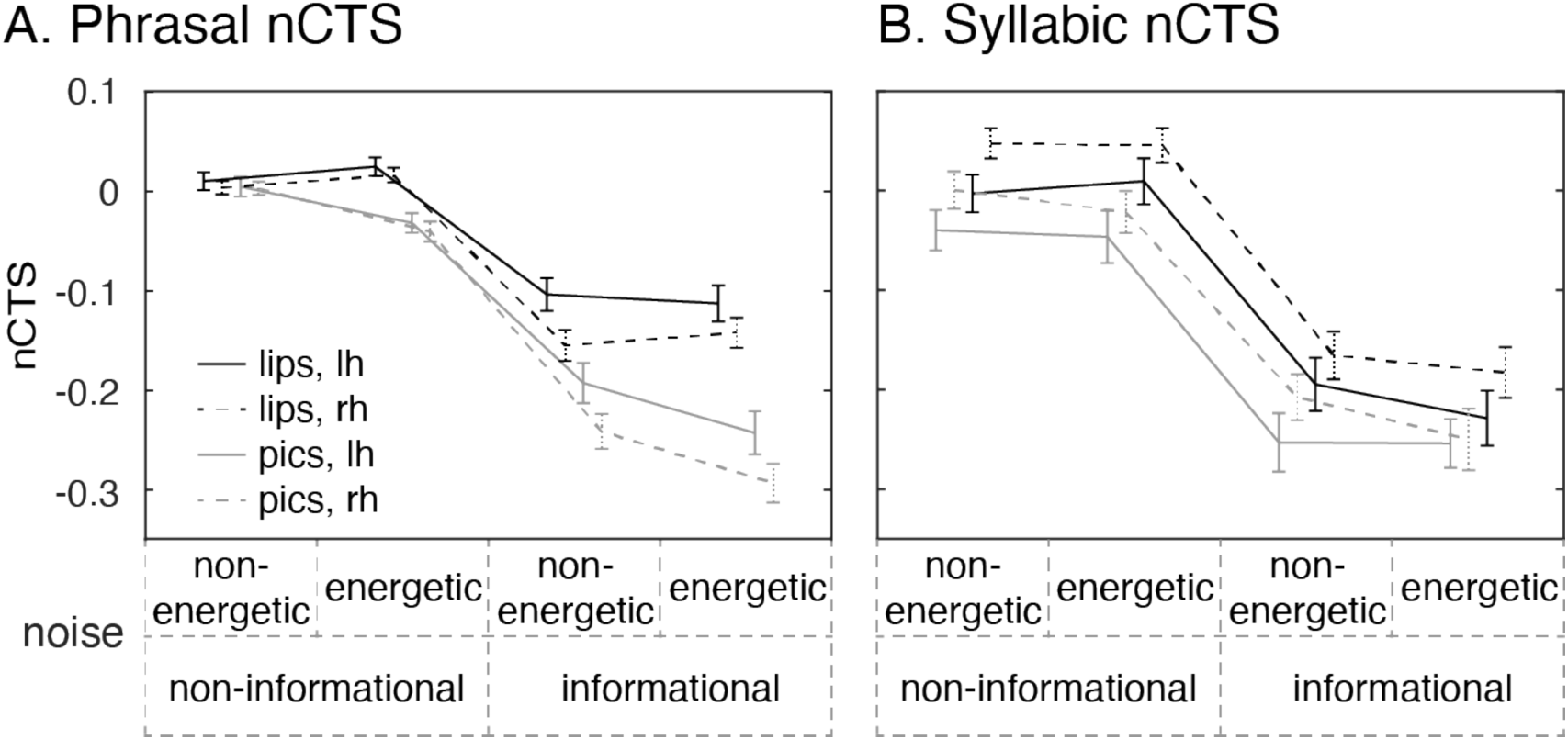
Impact of the main fixed-effects on the normalized cortical tracking of speech (nCTS) at phrasal (**A**) and syllabic rates (**B**). Mean and SEM values are displayed as a function of noise properties. The four traces correspond to visual conditions with the speaker’s talking face visible (*lips*; black traces) and with static pictures illustrating the story (*pics*; gray traces), within the left (*lh*; connected traces) and right (*rh*; dashed traces) hemispheres. nCTS values are bounded between –1 and 1, with values below 0 indicating lower CTS in speech-in-noise conditions than in noiseless conditions.

**Table 3.** Factors included in the final linear mixed-effects model fit to the normalized cortical tracking of speech (nCTS) at phrasal and syllabic rates. Factors are listed in their order of inclusion.

| | $\chi^2$ | | p |
| --- | --- | --- | --- |
|  | df | value |  |
| phrasal nCTS |  |  |  |
| noise | 3 | 597.50 | < 0.0001 |
| visual | 1 | 127.28 | < 0.0001 |
| hemisphere | 1 | 17.27 | < 0.0001 |
| noise × visual | 3 | 67.70 | < 0.0001 |
| noise × hemisphere | 3 | 10.98 | 0.012 |
| syllabic nCTS |  |  |  |
| noise | 3 | 340.61 | < 0.0001 |
| visual | 1 | 21.42 | < 0.0001 |
| hemisphere | 1 | 10.38 | 0.0013 |

The pattern of how nCTS changed with different types of noises was overall similar for phrasal and syllabic nCTS. Non-speech noise did not substantially change CTS (nCTS was close to 0). However, babble noise resulted in a substantial reduction of CTS compared to the noiseless condition for both hemispheres and irrespective of the availability of visual speech information. That is, nCTS in babble noise conditions was roughly between –0.1 and –0.3, indicating that CTS in babble noise was 20–50% (values obtained by inverting the formula of nCTS) lower than CTS in noiseless conditions.

Availability of visual speech information (*lips* conditions) increased the level of nCTS only in babble noise conditions for phrasal nCTS, and in all noise conditions for syllabic nCTS.

And finally, the noise impacted differently nCTS in the left and right hemispheres. The phrasal nCTS was higher in the left than right hemisphere in babble noise conditions. It was the other way round for syllabic nCTS in all noise conditions.

In summary, the CTS is mostly impacted by informational noises, and is also modulated by the availability of visual speech and the hemisphere (only in informational noise conditions for phrasal CTS, and in all noise conditions for syllabic CTS). These observations guided the elaboration of 8 relevant features (contrasts) of nCTS in SiN conditions (see Supplementary Methods): the global level of nCTS and its informational, visual and hemispheric modulations, all for phrasal and syllabic nCTS. In the next sections we unravel the associations between these features, reading abilities, and classical behavioral predictors of reading.

### What is the nature of the information about reading abilities brought by measures of SiN processing and classical behavioral predictors of reading?

Having identified relevant features of cortical SiN processing, we first evaluated to which extent these features and classical behavioral predictors of reading bring information about reading abilities. More precisely, we used partial information decomposition (PID) to dissect the information about reading abilities (target) brought by behavioral scores (first set of explanatory variables) and features of the nCTS in noise (second set of explanatory variables).^62–64^ Generally speaking, PID can reveal to which extent two sets of explanatory variables bring unique information about a target (information present in one set but not in the other), redundant information (information common to the two sets), and synergistic information (information emerging from the interaction of the two sets). Here, the target consisted of 5 reading scores: (i) an accuracy and (ii) a speed score for the reading of a connected meaningless text (Alouette test), and scores (number of correctly read words per unit of time) for the reading of a list of (iii) irregular words, (iv) regular words and (v) pseudowords. The first set of explanatory variables consisted of a total of 5 measures indexing phonological awareness (scores on phoneme suppression and fusion tasks), phonological memory (scores on forward and backward digit repetition), and lexical access (rapid automatized naming (RAN) score). The second set of explanatory variables was the 8 features of nCTS in SiN conditions identified in the previous subsection. Again, in that analysis, all measures were corrected for age, time spent at school and IQ.

As a result, features of nCTS in noise brought significant unique information about reading abilities (unique information = 0.61; *p* = 0.016), while classical behavioral predictors did not (unique information = 0.31; *p* = 0.10). Both sets of explanatory variables brought significant redundant but not synergistic information about reading (redundant information = 0.16; *p* = 0.0020; synergistic information = 0.12; *p* = 0.26).

These results indicate that the way the CTS is impacted by ambient noise relates to reading abilities in a way that is not fully explained by classical behavioral predictors of reading. Further analyses will therefore strive to identify which aspects of SiN processing and reading are related, and which of these relations are mediated by classical behavioral predictors of reading.

### Which features of SiN processing relate to reading abilities in a way that is not mediated by classical behavioral predictors of reading?

We next identified with linear mixed-effects modeling (i) the set of classical behavioral predictors of reading that best explains reading abilities, and (ii) the set of features of nCTS in noise that brings additional information about reading abilities. Importantly, all measures were corrected for age, time spent at school, and IQ, and further standardized. In that analysis, the type of reading score used to assess reading abilities was taken as a factor. Classical behavioral predictors of reading (5 measures) were first entered as regressors, before considering the features of nCTS in noise (8 measures) as additional regressors.

Table 4 presents the final linear mixed-effects model fit to reading scores. It shows that lexical access (indexed by RAN score) and phonological memory (indexed by the forward digit span) relate to global reading abilities. It also shows that two aspects of SiN processing, the visual and informational modulations in phrasal nCTS, explain a different part of the variance in reading abilities. Importantly, these two indices relate to reading in a way that depend on the type of reading score. These effects are illustrated with simple Pearson correlations in Table 5. The time necessary to fulfil the *RAN* task was significantly negatively correlated with all reading scores. The *forward digit span* was significantly positively correlated with all reading scores. The visual modulation in phrasal nCTS was overall positively correlated with scores involving reading speed (Alouette speed score and regular-, irregular- and pseudoword reading scores; significantly so for pseudoword reading only) but not with the Alouette accuracy score. The informational modulation in phrasal nCTS was characterized by a significant positive correlation with the score on irregular word reading only. Interestingly, the correlation was not significant—and even negative—with the score on pseudoword reading.

**Table 4.** Regressors included in the final linear mixed-effects model fit to the 5 reading scores taken as factors. Regressors are listed in their order of inclusion.

| | $\chi^2$ | | p |
| --- | --- | --- | --- |
|  | df | value |  |
| RAN | 1 | 15.79 | < 0.0001 |
| forward digit span | 1 | 11.10 | 0.0009 |
| visual modulation in phrasal nCTS | 1 | 4.85 | 0.028 |
| informational modulation in phrasal nCTS dependant on reading score | 5 | 15.63 | 0.0080 |
| visual modulation in phrasal nCTS dependant on reading score | 4 | 11.09 | 0.026 |

**Table 5.** Pearson correlation between measures of reading abilities and relevant brain and behavioral measures. *** *p* < 0.001, ** *p* < 0.01, * *p* < 0.05, ^#^ *p* < 0.1. nCTS, normalized cortical tracking of speech.

|  | RAN | forward digit span | Visual modulation in phrasal nCTS | Informational modulation in phrasal nCTS | Visual modulation in syllabic nCTS | Phoneme suppression | Phoneme fusion |
| --- | --- | --- | --- | --- | --- | --- | --- |
| Alouette accuracy | **-0.37 \*\*** | **0.33 \*\*** | 0.00 | -0.35 | **0.29 \*** | 0.11 | **0.25 \*** |
| Alouette speed | **-0.41 \*\*\*** | **0.38 \*\*\*** | 0.21 # | 0.08 | **0.30 \*\*** | **0.31 \*\*** | **0.30 \*\*** |
| Irregular words | **-0.35 \*\*** | **0.42 \*\*\*** | 0.18 | **0.26 \*** | **0.37 \*\*** | 0.21 # | 0.17 |
| Regular words | **-0.42 \*\*\*** | **0.35 \*\*** | 0.18 | 0.12 | **0.31 \*\*** | **0.25 \*** | 0.19 |
| Pseudowords | **-0.34 \*\*** | **0.30 \*** | **0.31 \*\*** | -0.07 | **0.23 \*** | 0.21 # | 0.11 |

We will now attempt to better understand the meaning of this last association (between the informational modulation in phrasal nCTS and irregular- but not pseudoword reading). Given that different routes support reading of irregular words (lexical route) and pseudowords (sublexical route), the contrast between corresponding standardized scores (irregular – pseudowords) indicates reading strategy. We henceforth refer to this index as the reading strategy index. Further strengthening the correlation pattern highlighted above for the informational modulation in phrasal nCTS, this latter index correlated even more strongly with the reading strategy index (*r* = 0.44, *p* < 0.0001; See Fig. 3, left) than with the score on irregular word reading. This suggests that irregular and pseudoword reading scores bring synergistic information about the informational modulation in phrasal nCTS. To confirm this, we used partial information decomposition (PID) to dissect the information about the informational modulation in phrasal nCTS (target) brought by irregular reading scores (first explanatory variable) and pseudoword reading scores (second explanatory variable). This analysis revealed that the score on irregular word reading carried significant unique information about the informational modulation in phrasal nCTS (unique information = 0.044, *p* = 0.015), while the score on pseudowords did not (unique information = 0.0001, *p* = 0.62), and most interestingly, that these two reading scores carried significant synergistic but not redundant information about the informational modulation in phrasal nCTS (redundant information = 0.0019, *p* = 0.44; synergistic information = 0.146, *p* < 0.0001).

**Figure 3.**
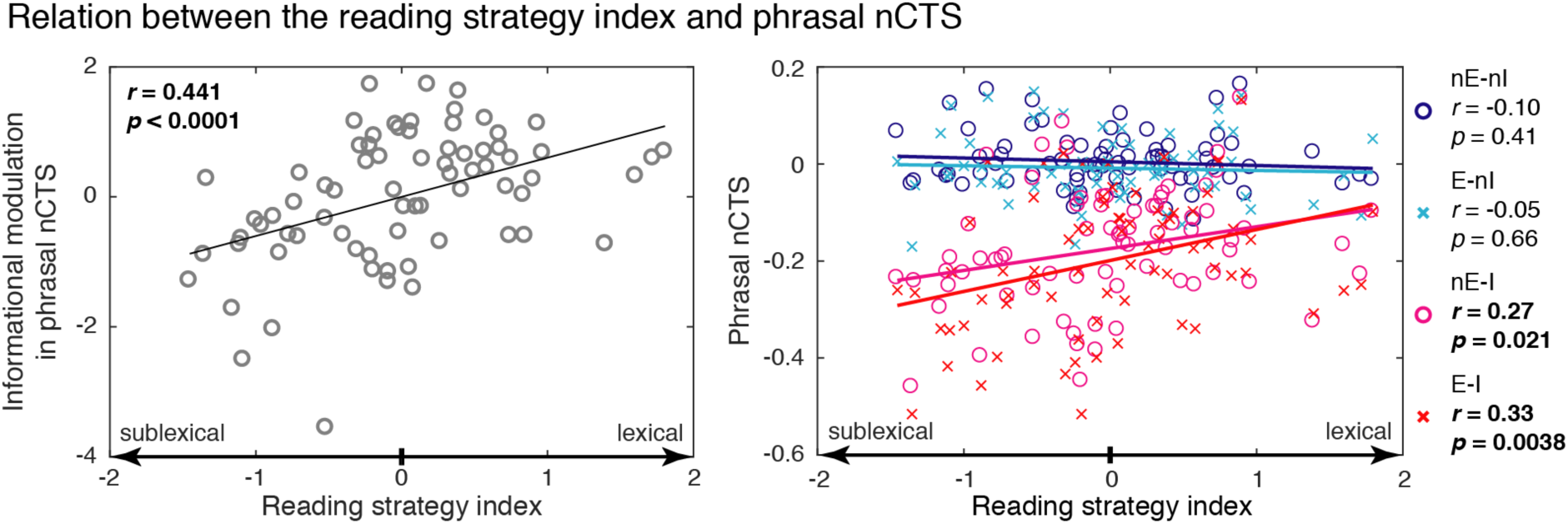
Relation between the reading strategy index and the normalized cortical tracking of speech (nCTS) at phrasal rate. **Left** — The informational modulation in phrasal nCTS as function of the reading strategy index. Gray circles depict participants’ values and a black trace is the regression line, with correlation and significance values indicated in the bottom right corner. **Right** — The mean nCTS across visual conditions and both hemispheres for the 4 types of noise: energetic (*E*) or not (*nE*) and informational (*I*) or not (*nI*), with *nE-nI* in blue, *E-nI* in turquoise, *nE-I* in red and *E-I* in pink. Circles (*nE* noise conditions) and crosses (*E*) depict participants’ values, and full traces are the regression lines. Correlation and significance level for all noise conditions are indicated on the right of each plot.

Figure 3 (right panel) further illustrates that the reading strategy index was correlated with phrasal nCTS only in the informational noise conditions.

In summary, classical behavioral predictors of reading were informative about global reading abilities (similar correlation with all 5 measures of reading), while two aspects of the CTS in noise (informational and visual modulations in phrasal nCTS) related to specific aspects of reading (correlation with some but not all 5 measures of reading). The extent to which visual speech boosts phrasal CTS in noise was related to reading speed but not accuracy, and the ability to maintain adequate phrasal CTS in babble noise related to reading strategy (dominant reliance on the lexical rather than sublexical route).

### Do other features of SiN processing or classical behavioral predictors of reading relate to reading abilities?

Above, we have identified a set of brain and behavioral measures related to reading. Importantly, each measure was included because it explained a new part of the variance in reading abilities. But the first PID analysis revealed that brain and behavioral measures do carry significant redundant information. This means that some measures might have been left aside if they explained some variance that was already explained (i.e., if they provided mainly redundant information). Accordingly, we also ran the linear mixed-effects analysis with nCTS and behavioral regressors that were not included. This analysis identified an overall positive correlation between reading abilities and (i) the visual modulation in syllabic nCTS (*χ*^2^(1) = 9.74, *p* = 0.0018), (ii) phoneme suppression (*χ*^2^(1) = 4.94, *p* = 0.026) and (iii) phoneme fusion (*χ*^2^(1) = 4.00, *p* = 0.038). Corresponding Pearson correlation coefficients are presented in Table 5. A detailed PID analysis revealed that these “side” measures were redundant—and synergistic to some extent—with RAN and forward digit span but not with visual and informational modulations in phrasal nCTS (see Supplementary Results).

In summary, scores indexing phonological awareness (score on phoneme suppression and phoneme fusion) and the extent to which visual speech boosts syllabic CTS in noise (visual modulation in syllabic nCTS) relate to global reading abilities in a way that is mediated by the main classical behavioral predictors of reading we identified (RAN and forward digit span) but not with visual and informational modulations in phrasal nCTS

### Does phonological awareness mediate SiN perception capacities?

Having identified three relations between various aspects of cortical SiN processing and reading, we now specifically test the hypothesis that each of these relations is mediated by phonological awareness. For that, we again relied on PID to decompose the information about reading abilities (target) brought by each identified feature of the CTS in noise (first explanatory variable) and the mean of the two scores indexing phonological awareness (second explanatory variable). Ensuing results are provided in Table S1. In summary, phonological awareness mediated one aspect of the relation between reading and cortical SiN processing (relation with the benefit of visual speech to boost syllabic CTS in noise), but not the two others (relations involving phrasal CTS in noise).

### Do relations between reading and features of nCTS translate to alterations in dyslexia?

We next evaluated whether the relations between features of nCTS and reading abilities translate to alterations in dyslexia. That analysis was conducted on a group of 26 dyslexic readers, 26 age-matched and 26 reading-level-matched typically developing children selected among the 73 children included in the first part of the study.

Based on the result that reading abilities relate to phrasal nCTS in informational noise and to the boost in nCTS brought by visual speech, we focused the comparison on the phrasal nCTS in *lips* and *pics* averaged across hemispheres and babble noise conditions (see Fig. 4A). As a result, individuals with dyslexia had significantly lower phrasal nCTS than age-matched controls in *pics* (*t*(50) = 3.03, *p* = 0.0039) but not in *lips* (*t*(50) = 0.83, *p* = 0.41). This difference was not present in the comparison with reading-level-matched controls (*pics*, *t*(50) = 0.65, *p* = 0.52; *lips*, *t*(50) = 0.54, *p* = 0.59).

**Figure 4.**
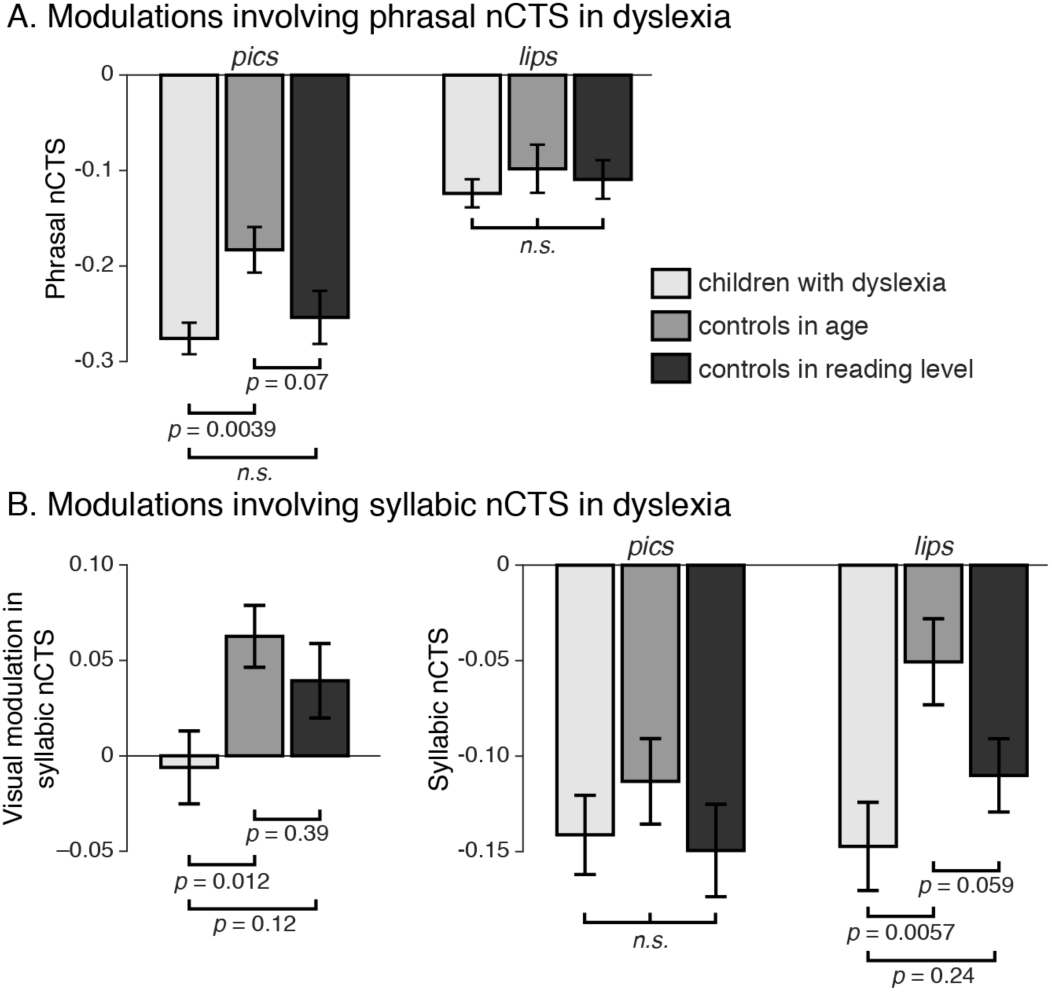
Comparison between children with dyslexia and controls in the measures of normalized cortical tracking of speech (nCTS) significantly related to reading abilities. **A** — Modulations involving phrasal nCTS. Displayed are the mean and SEM within groups (dyslexia, control in age, and control in reading level) of phrasal nCTS in the conditions with (*lips*) and without (*pics*) visual speech information. Values of nCTS were averaged across hemispheres and informational noise conditions for phrasal nCTS and across hemispheres and all noise conditions for syllabic nCTS. **B** — Modulations involving syllabic nCTS. On the left is the visual modulation in syllabic nCTS. The right part is as in A.

Based on the result that reading abilities relate to the visual modulation in syllabic nCTS, we focused the comparison on this index (see Fig. 4B left part). This revealed that individuals with dyslexia had significantly lower visual modulation in syllabic nCTS than age-matched (*t*(50) = 2.62, *p* = 0.012) but not reading-level-matched controls (*t*(50) = 1.59, *p* = 0.12). To better understand the nature of this difference, we further compared between groups the syllabic nCTS in *lips* and *pics* averaged across hemispheres and noise conditions (see Fig. 4B right part). As a result, individuals with dyslexia had significantly lower syllabic nCTS than age-matched controls in *lips* (*t*(50) = 2.89, *p* = 0.0057) but not in *pics* (*t*(50) = 0.88, *p* = 0.38). This difference was not present in the comparison with reading-level-matched controls (*lips*, *t*(50) = 1.19, *p* = 0.24; *pics*, *t*(50) = 0.25, *p* = 0.81).

In summary, one aspect of cortical SiN processing (reliance on visual speech to boost phrasal nCTS) was not altered in dyslexia while two other aspects (phrasal nCTS in babble noise, and reliance on visual speech to boost syllabic nCTS) were altered in dyslexia in comparison with typical readers matched for age but not reading level. This suggests that these two later aspects are altered as a consequence of reduced reading experience.

## Discussion

The main objective of this study was to fully characterize the nature of the relation between objective cortical measures of SiN processing and reading abilities in elementary-school children. Results demonstrate that some cortical measures of SiN processing relate to reading performance and reading strategy. First, phrasal nCTS in babble (*i.e.*, informational) noise relates to the ability to read irregular but not pseudowords, which in the DRC model indicates maturation of the lexical route. Second, the ability to leverage visual speech to boost phrasal nCTS in babble noise relates to reading speed (but not accuracy). Third, the ability to leverage visual speech to boost syllabic nCTS in noise relates to global reading abilities. Fourth, classical behavioral predictors of reading abilities (RAN, phonological memory and phonological awareness) relate to global reading performance, and not strategy. Importantly, behavioral scores and the two features of phrasal CTS in babble noise explained a different part of the variance in reading abilities. Finally, the first and third relations uncovered in typical readers translated to significant alteration in dyslexia in comparison with aged-matched but not reading-level-matched typically developing children. Two limitations are discussed in Supplementary Discussion.

Significant associations were found between reading abilities and some features of phrasal and syllabic nCTS. There is evidence that CTS at phrasal rate (here taken as 0.2–1.5 Hz) partly reflects parsing or chunking of words, phrases and sentences.^65^ Indeed, the brain tracks phrase and sentence boundaries even when speech is devoid of prosody but only if it is comprehensible;^39^ and the phase of below-4-Hz brain oscillations modulates perception of ambiguous sentences.^37^ CTS at phrasal/sentential rate would help align neural excitability with syntactic information to optimize language comprehension.^36^ In contrast, CTS at syllable rate (here taken as 2–8 Hz) would reflect low level auditory processing.^65^ In light of the above, our results highlight that associations between SiN perception and reading abilities build on their shared reliance on both language processing and low-level auditory processing.

### Robustness of cortical speech representation to babble noise indexes the development of the lexical route

Our results indicate that an objective cortical measure of the ability to deal with babble noise relates to the maturation of the lexical route. Technically, the informational modulation in phrasal nCTS correlated significantly positively with the reading score on irregular but not pseudowords. Reading score on irregular words indeed provided unique information about the informational modulation in nCTS. Also, the two reading scores in synergy provided some additional information about the informational modulation in nCTS. Furthermore, the result that the informational modulation in nCTS correlated more with the reading strategy index than the score on the irregular words suggests that the key elements at the basis of this relation are the processes needed to read irregular words that are not needed to read pseudo-words. Our results in dyslexia support this relation although they cannot rule out the possibility that it is due to variability in reading experience since phrasal nCTS in non-visual informational noise conditions was reduced in dyslexic readers compared with age-matched but not reading-level-matched controls.

The relation between the development of the lexical route and the level of phrasal nCTS in babble noise could be explained by a positive influence of good SiN abilities on reading acquisition. Let us take as an example the situation of being faced for the first time with a written word that is read by a teacher while some classmates are making noise. SiN abilities will naturally determine the odds of hearing that word properly and hence the odds of building up the orthographic lexicon. When reading again the word alone, only children with good SiN abilities will have the opportunity to train their lexical route for that specific word. Of course the same chain of action could be posited for the training of grapheme–phoneme correspondence. But there are many more words than phonemes and syllables, so that good SiN abilities might be more important to successfully learn the correspondence between irregular words’ orthographic and phonological representations. Indeed, grapheme–phoneme correspondence is intensively trained when learning to read. Children are repeatedly exposed to examples of successful grapheme–phoneme correspondence, some with noise, and some without noise. Accordingly, no matter what children’s SiN abilities are, they will learn the grapheme–phoneme correspondence and develop their sublexical route, provided that they have adequate phonological awareness. Supporting this, phonological awareness does not predict SiN abilities in typical readers.^21^

Alternatively, the relation between the ability to read irregular words (which tags the development of the lexical route) and nCTS in babble noise could be mediated by the degree of maturation of the mental lexicon.^66,67^ The mental lexicon integrates and binds the orthographic, semantic and phonological representations of words. Its proper development is important for reading acquisition. Indeed, reading acquisition entails creating a new orthographic lexicon and binding it to the preexisting semantic and phonological lexicons.^68^ Development of such binding (i) is indispensable to read irregular words,^69^ (ii) benefits reading of regular words, and (iii) does not contribute to reading pseudowords. The proper development of the mental lexicon is also important for SiN comprehension. Indeed, SiN comprehension strongly depends on vocabulary knowledge.^21,70,71^ And the level of CTS in noise relates to the listeners’ level of comprehension.^35,40,41^ This therefore suggests that the robustness of CTS to babble noise depends on the level of comprehension, which in turn depends on how developed is the mental lexicon. The development of the mental lexicon could therefore be the hidden factor mediating the relation between SiN and lexical reading ability. This is also perfectly in line with our result that altered phrasal nCTS in babble noise in dyslexia may result from reduced reading experience. In brief, reading difficulties in dyslexia would reduce their reading experience, which would impair building up the mental lexicon, and in turn impede SiN perception. Still, future studies on the association between SiN processing and reading should include measures of the development of the mental lexicon to carefully analyse the interrelation between SiN perception, reading abilities and the development of the mental lexicon.

### Audiovisual integration and reading abilities

We found significant relations between reading abilities and the ability to leverage visual speech to maintain phrasal and syllabic CTS in noise. Visual speech cues (articulatory mouth and facial gestures) are well known to benefit SiN comprehension ^55^ and CTS in noise.^72–76^ Obviously, the auditory signal carries much more fine-grained information about the phonemic content of speech than the visual signal. But the effect of audiovisual speech integration is quite evident in SiN conditions, where it affords a substantial comprehension benefit.^55,56,77,78^ Mirroring this perceptual benefit, it is already well documented that phrasal and syllabic CTS in noise is boosted in adults when visual speech information is available.^72–76,79–82^

We found that the visual modulation in phrasal nCTS correlated globally positively with reading speed (significantly so for the pseudowords) but not accuracy. However, our dyslexic readers (compared with both control groups) did not have any alteration in their phrasal nCTS in babble noise when visual speech was provided. Instead, they successfully relied on visual speech information to restore their phrasal CTS in babble noise (which was altered without visual speech information). In other words, reliance on lip-reading to maintain appropriate phrasal CTS in babble noise appeared as a protection factor in our group of dyslexic readers.

We also found that the visual modulation in syllabic nCTS correlated globally positively with reading abilities. More interestingly, our dyslexic readers (compared with both control groups) did not have any significant alteration in their syllabic nCTS in noise when visual speech was not provided. However, compared with age-matched typically developing children, they benefited significantly less from visual speech to boost syllabic CTS in noise. Instead, they behaved more like reading-level-matched typically developing children. Accordingly, our results cannot argue against the view that poor audiovisual integration in dyslexia is caused by reduced reading experience.^57,83,84^ Notwithstanding, the pattern of results (see Fig. 4B left) is even suggestive of an alteration in dyslexia in comparison with reading-level-matched children. More statistical power would be needed to confirm/infirm the trend.

Our result that audiovisual integration abilities correlate with reading abilities is in line with existing literature. Indeed, individuals with dyslexia benefit less from visual cues to perceive SiN than typical readers^85–89^. Audiovisual integration and reading could be altered in dyslexia simply because both rely on similar mechanisms. Indeed, reading relies on the ability to bind visual (graphemic) and auditory (phonemic) speech representations.^90,91^ And according to some authors, suboptimal audiovisual integration mechanisms could reduce reading fluency.^92^ Importantly, the finding that individuals with dyslexia benefit normally from visual speech to boost phrasal but not syllabic CTS in noise brings important information about the nature of the audiovisual integration deficit in dyslexia. Following the functional roles attributed to CTS, individuals with dyslexia would properly integrate visual speech information to optimize processing of syntactic information,^36^ but not to support acoustic/phonemic processing.^65^ This could be explained by their preserved ability to extract and integrate the temporal dynamics of visual speech, but not the lip configuration,^89^ two aspects of audiovisual speech integration currently thought to be supported by distinct neuronal pathways.^93^ This inability to rely on lip configuration to improve auditory phonemic perception in SiN conditions may be caused by a supra-modal phonemic categorization deficit, as already proposed for children with specific language impairment.^94^ Finally, the fact that the visual modulation in syllabic nCTS brought a limited amount of unique information about reading with respect to classical behavioral predictors of reading, but that all of them brought more information in synergy, suggests that a broad set of low-level processing abilities contribute to determine reading abilities and alterations in dyslexia.^95,96^

### Classical behavioral predictors related to global reading abilities

Our results confirm that classical behavioral predictors of reading (RAN, phonological memory, and metaphonological abilities) are directly related to the global reading level rather than reading strategy. We draw this conclusion since the optimal model for reading score contained a common slope for all reading subtests. This means that the model was not significantly improved by optimizing the slope for each of the 5 reading subtests separately. Accordingly, univariate correlation coefficients presented in Table 5 were roughly similar across the 5 reading scores.

Phonological memory (assessed with forward digit span) was significantly positively correlated with the global reading level. That phonological memory relates to global reading abilities rather than reading strategy is well documented.^4^ Poor readers, regardless of their reading profile, typically perform poorly on phonological memory tests involving digits, letters,^97,98^ or words.^99^

Performance on the RAN task was also related to the global reading level, in line with existing literature.^6–8,100–103^ RAN performance has indeed a moderate-to-strong relationship with all classical reading measures alike, including word, non-word and text reading, as well as text comprehension.^100^ It is a consistent predictor of reading fluency in various alphabetic orthographies independently of their complexity.^104^ RAN performance even predicts reading performance at 2 years interval,^105^ similarly well for reading performance assessed with tasks tagging lexical and sublexical routes. It is thought that RAN and reading performances correlate because both involve serial processing and oral production,^103^ two processes that are common to both reading routes.

Finally, phonological awareness assessed with phoneme suppression and fusion tasks was significantly related to reading abilities. However, the information it brought about reading was less, and essentially redundant with that brought by RAN and phonological memory. This is not surprising given that children tested in the present study had at least one year of reading experience. Phonological awareness indeed plays a key role in the early stages of reading acquisition, i.e., when learning grapheme-to-phoneme conversion,^106–108^ and undergoes a substantial maturation during that period.^109^

### Phonological awareness

Our results indicate that, in typical readers, phonological awareness mediates at best part of the relation between the cortical processing of SiN and reading abilities. Indeed, the information about reading brought by phonological awareness was redundant with that brought by the visual modulation in syllabic nCTS, but not with that brought by the informational and visual modulations in phrasal nCTS. This finding illustrates the importance of separating the different processes involved in SiN processing and reading to seek associations. It also provides a potential reason why contradictory reports exist on the topic.^19–21^

## Conclusion

Overall, these results significantly further our understanding of the nature of the relation between SiN processing abilities and reading abilities. They demonstrate that cortical processing of SiN and reading abilities are related in several specific ways, and that some of these relations translate into alterations in dyslexia that are attributable to reading experience. They also demonstrate that classical behavioral predictors of reading (including phonological awareness) mediate relations involving the processing of acoustic/phonemic but not syntactic information in natural SiN conditions. This contrasts with the classically assumed mediating role of phonological awareness. Instead, the ability to process speech syntax in babble noise could directly modulate skilled reading acquisition. Finally, the information about reading abilities brought by cortical markers of syntactic processing of SiN was complementary to that provided by classical behavioral predictors of reading. This implies that such markers of SiN processing could serve as novel electrophysiological markers of reading abilities.

## Material and Methods

### Participants

Seventy-three typical and 26 dyslexic readers enrolled in elementary-school took part in this experiment (see Table 1 for participants’ characteristics). All were native French speakers, reported being right-handed, had normal hearing according to pure tone audiometry (normal hearing thresholds between 0–25 dB HL for 250, 500, 1000, 2000, 4000 and 8000 Hz) and normal SiN perception as revealed by a SiN test (Lafon 30) from a French language central auditory battery.^110^ We used a French translation of the Family Affluence Scale ^111^ to evaluate participants’ socio-economic level.

This study was approved by the Ethics Committee of the CUB Hôpital Erasme (Brussels, Belgium). Participants and their legal representatives signed a written informed consent before participation. Participants were compensated with a gift card worth 50 euros.

### Behavioral assessment

Participants underwent a comprehensive behavioral assessment intended to appraise their reading abilities and some cognitive abilities related to reading or speech perception.

#### Reading abilities

Children completed the word reading (regular, irregular and pseudowords) tasks of a dyslexia detection tool (ODEDYS-2; ^112^ and the Alouette-R reading task ^113^).

For each of the word reading tasks (regular, irregular or pseudowords), participants had to read as rapidly and accurately as possible a list of 20 words. Each task provided a *reading* score computed as the number of words correctly read divided by the reading time (in seconds).

In the Alouette-R task,^113^ children had 3 minutes to read as rapidly and accurately as possible a text of 256 words. This text is composed by a succession of words which do not tell a meaningful story. This peculiarity forces children to solely rely on their reading skills and prevents children from using anticipation or inference strategies that could boost the reading scores. An *accuracy* score was computed as the number of words correctly read divided by the total number of words read, and a *speed* score as the number of words correctly read multiplied by the ratio of 180 s (maximal reading time) to the effective reading time.

#### Phonological processing

The initial phoneme suppression and initial phonemes fusion tasks of the ODEDYS-2^112^ was used to assess phonological processing.

In the initial phoneme suppression task, children had to repeat orally presented words while intentionally suppressing the initial phoneme of the word (*i.e.* dog -> og). In total, 10 words were presented, and performance was quantified as percentage correct.

In the initial phoneme fusion task, children had to combine the initial phoneme of two orally presented words to create a new (non-)word (*i.e., Big & Owen -> /bo/*). In total, 10 pairs of words were presented, and performance was quantified as percentage correct.

#### Rapid Automatized Naming

We used the RAN task of the ODEDYS-2.^112^ Children had to name as rapidly and accurately as possible 25 pictures (5 different pictures randomly repeated 5 times). Performance was quantified as the total time to complete the task, meaning that the lower the score, the better the performance.

#### Phonological memory

The forward and backward digit repetition task from the ODEDYS-2 ^112^ was used to assess phonological memory.

In the forward digit repetition task, children were asked to repeat orally presented numbers series in the same order as presented. The series are different at every trial. The first series contains 3 digits, and the size of the series is incremented by one every second trial. The task ends after a failure to repeat the 2 series of a given size. Forward digit span score was taken as the number of digits in the last correctly repeated series.

The backward digit repetition task is akin to the forward one. The only difference is that digit series have to be repeated in the exact reverse order (e.g., children presented 1 2 3 4 have to repeat 4 3 2 1).

#### Attention abilities

The Bells test ^114^ was used to assess visual attention and the TAP auditory attention subtest ^115^ to assess the auditory attentional level.

In the Bells test, children had 2 minutes to find as many bells as possible on a sheet comprising 35 bells scattered among 280 visual distractors. Performance was quantified as the number of bells found divided by the time needed.

In the TAP auditory attention subtest, a, children had to focus their attention during 3 min 20 s on an auditory stream. Children were hearing a train of 200 pure tone stimuli lasting 500 ms with a 1000-ms stimulus onset asynchrony. Tones alternated between high (1073 Hz) and low (450 Hz) pitch. There were 16 occurrences in which 2 high or low pitch tones were following one another. Only in this case, participants had to press a response button as fast as possible. A performance score was quantified as the number of correct responses, a speed score as the mean response time, and a failure score as the number of responses to tones differing in pitch with the preceding one.

#### Non-verbal intelligence

The brief version of the Weschler Nonverbal (WNV) Scale of Ability ^116^ was used to assess non-verbal intelligence.

This assessment consisted of matrices and recognition subtests for children younger than 8 years. Older children were assessed with matrices and spatial memory subtests.

In matrices subtest, children were presented with incomplete visual matrices and had to select the good missing portion among 4 or 5 response options. The subtest ended when 4 mistakes were made in the last 5 trials. A raw score was taken as the number of correctly completed matrices. This raw score was converted to a *T* score by comparison with values provided in a table of norms.

In recognition subtest, children had to carefully look at visual geometric designs that were presented one by one for three seconds. After each presentation, they had to identify the previously seen design among four or five response options. The subtest ended when 4 mistakes were made in the last 5 recognition trials. A raw score was taken as the number of correctly recognized drawings. This raw score was converted to a *T* score by comparison with values provided in a table of norms.

In spatial memory subtest, children were presented with a board with 10 cubes spread on it, and were asked to mimic the examiner’s tapping sequence. The sequences are different on every trial. The first sequence consists in tapping on 2 cubes, and the size of the sequences is incremented by one every second trial. The task ends after a failure to repeat 2 sequences of a given size. This task was performed twice, in the forward and backward directions. For each direction, a raw score was taken as the number of correctly repeated sequences. Raw scores were summed and converted to a *T* score by comparison with values provided in a table of norms.

Total non-verbal IQ was computed as the sum of both *T* scores, which compared to a table of norms provided a *Total non-verbal* IQ score.

### Neuroimaging assessment

#### Stimuli

The stimuli were derived from 12 audiovisual recordings of 4 native French-speaking actors (2 females, 3 recordings per actor) telling a story for ~6-min (mean ± SD, 6.0 ± 0.8 min) (see Supplementary Methods for details on recording of video stimuli). In each video, the first 5 s were kept unaltered to enable children to unambiguously identify the actor’s voice and face they were requested to attend to. The remainder of the video was divided into 10 consecutive blocks of equal size that were assigned to 9 conditions. Two blocks were assigned to the *noiseless* condition in which the audio track was kept but the video was replaced by static pictures illustrating the story (mean ± SD picture presentation time across all videos, 27.7 ± 10.8 s). The remaining 8 blocks were assigned to 8 conditions in which the original sound was mixed with a background noise at 3 dB signal-to-noise ratio (SNR). There were 4 different types of noise, and each type of noise was presented once with the original video, thereby giving access to lip-read information (*lips* visual conditions), and once with the static pictures illustrating the story (*pics* visual conditions). The different types of noise differed in the degree of energetic and informational interference they introduced.^54^ The non-energetic non-informational noise was a white noise filtered through 100–10000-Hz. The (maximally-)energetic non-informational noise had its spectral properties dynamically adapted to mirror those of the actor’s voice ~1 s around (see Supplementary Methods for the procedure used to build the energetic non-informational noise). The non-(or least-)energetic informational noise was a 5-talker cocktail party noise recorded by individuals of gender opposite to the actor’s (i.e., a 5-man for female actors). The (maximally-)energetic informational noise was a 5-talker cocktail party noise recorded by individuals of gender identical to the actor’s. The assignment of conditions to blocks was random, with the constraint that each of the 5 first and last blocks contained exactly 1 *noiseless* audio, 2 *lips* videos, 2 energetic noises, and 2 informational noises. Smooth audio and video transitions between blocks was ensured with 2-s fade-in and fade-out. Ensuing videos were grouped in 3 disjoint sets featuring one video of each of the actors (total set duration: 23.0, 24.3, 24.65 min), and there were 4 versions of each set differing in condition random ordering.

#### Experimental paradigm

During the imaging session, participants were laying on a bed with their head inside the MEG helmet. Their brain activity was recorded while they were attending 4 videos (separate recording for each video) of a randomly selected set and ordering of the videos presented in a random order, and finally while they were at *rest* (eyes opened, fixation cross) for 5 min. They were instructed to watch the videos attentively, listen to the actors’ voice while ignoring the interfering noise, and remain as still as possible. After each video, they were asked 10 yes/no comprehension questions related to each of the 10 blocks/conditions (data not analyzed here). Videos were projected onto a back-projection screen placed vertically, ~120 cm away from the MEG helmet. The inner dimensions of the black frame were 35.2 cm (horizontal) and 28.8 cm (vertical), and actors face spanned ~15 cm (horizontal) and ~20 cm (vertical). Participants could see the screen through a mirror placed above their head. In total the optical path from the screen to participants’ eyes was of ~150 cm. Sounds were delivered at 60 dB (measured at ear-level) through a MEG-compatible front-facing flat-panel loudspeaker (Panphonics Oy, Espoo, Finland) placed ~1 m behind the screen.

#### Data acquisition

During the experimental conditions, participants’ brain activity was recorded with MEG at the CUB Hôpital Erasme. Neuromagnetic signals were recorded with a whole-scalp-covering MEG system (Triux, Elekta) placed in a lightweight magnetically shielded room (Maxshield, Elekta), the characteristics of which being described elsewhere.^117^ The sensor array of the MEG system comprised 306 sensors arranged in 102 triplets of one magnetometer and two orthogonal planar gradiometers. Magnetometers measure the radial component of the magnetic field, while planar gradiometers measure its spatial derivative in the tangential directions. MEG signals were band-pass filtered at 0.1–330 Hz and sampled at 1000 Hz.

We used 4 head-position indicator coils to monitor subjects’ head position during the experimentation. Before the MEG session, we digitized the location of these coils and at least 300 head-surface points (on scalp, nose, and face) with respect to anatomical fiducials with an electromagnetic tracker (Fastrack, Polhemus).

Finally, subjects’ high-resolution 3D-T1 cerebral images were acquired with a magnetic resonance imaging (MRI) scanner (MRI 1.5T, Intera, Philips) after the MEG session.

#### Data pre-processing

Continuous MEG data were first preprocessed off-line using the temporal signal space separation method implemented in MaxFilter software (MaxFilter, Neuromag, Elekta; correlation limit 0.9, segment length 20 s) to suppress external interferences and to correct for head movements.^118,119^ To further suppress physiological artifacts, 30 independent components were evaluated from the data band-pass filtered at 0.1–25 Hz and reduced to a rank of 30 with principal component analysis. Independent components corresponding to heartbeat, eye-blink, and eye-movement artifacts were identified, and corresponding MEG signals reconstructed by means of the mixing matrix were subtracted from the full-rank data. Across subjects and conditions, the number of subtracted components was 3.45 ± 1.23 (mean ± SD across subjects and recordings). Finally, a window time of 1 s time points at timings 1 s around remaining artifacts were set to bad. Data were considered contaminated by artifacts when MEG amplitude exceeded 5 pT in at least one magnetometer or 1 pT/cm in at least one gradiometer.

We extracted the temporal envelope of the attended speech (actors’ voice) using the optimal approach proposed by Biesmans et al.^120^. Briefly, audio signals were bandpass filtered using a gammatone filter bank (15 filters centered on logarithmically-spaced frequencies from 150 Hz to 4000 Hz), and subband envelopes were computed using Hilbert transform, elevated to the power 0.6, and averaged across bands.

#### Accuracy of speech envelope reconstruction and normalized CTS

For each condition and participant, a global value of cortical tracking of the attended speech was evaluated for all left-hemisphere sensors at once, and for all right-hemisphere sensors at once. Using the mTRF toolbox,^58^ we trained a decoder on MEG data to reconstruct speech temporal envelope, and estimated its Pearson correlation with real speech temporal envelope. This correlation is often referred to as the reconstruction accuracy, and it provides a global measure of cortical tracking of speech. See Supplementary Methods for a full description of the procedure and statistical assessment. A similar approach has been used in previous studies on the cortical tracking of speech.^47,51,60,61^

Based on CTS values in noiseless condition (CTS_noiseless_) and in each SiN condition (CTS_SiN_), we estimated nCTS as follows:

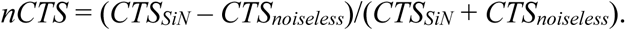

This index can however be misleading when derived from negative CTS values (which may happen since CTS is an unsquared correlation values). For this reason, and for the sake of nCTS computation only, CTS values below a threshold of 10% of the mean CTS across all subjects, conditions and hemispheres were set to that threshold prior to nCTS computation.

### Linear mixed-effects modeling of nCTS and reading values

All behavioral and nCTS measures were corrected for IQ, age, time spent at elementary school, and for outliers (see Supplementary Methods for details on this procedure).

We performed linear mixed-effects analysis with R ^121^ and *lme4* ^122^ to identify how different fixed effects modulate nCTS. We started with a null model that included only a different random intercept for each subject. The model was iteratively compared to models incremented with simple fixed effects of *hemisphere*, *noise* (non-energetic non-informational, energetic non-informational, non-energetic informational, and energetic informational), and *visual* (*lips* vs. *pics*) added one by one. At every step, the most significant fixed effect was retained, until the addition of the remaining effects did not improve the model any further (*p* > 0.05). The same procedure was then repeated to refine the ensuing model with the interactions of the simple fixed effects of order 2 (e.g., *hemisphere* × *noise*) and then 3 (*hemisphere* × *noise* × *visual*).

We followed the same approach to identify how reading abilities (5 standardized scores) relate to classical behavioral predictors of reading and features of nCTS. In that analysis, we first considered a non-zero slope for the classical behavioral predictors identical for all reading scores, then a non-zero slope for the classical behavioral predictors different for all reading scores, then a non-zero slope for the features of nCTS identical for all reading scores, and finally a non-zero slope for the features of nCTS different for all reading scores.

### Partial information decomposition

We used PID to appraise without *a priori* the relation between reading abilities, cortical measures of SiN processing, and classical behavioral predictors of reading. In general, PID decomposes the mutual information (MI) quantifying the relationship between two explanatory variables (or sets of explanatory variables) and a single target, into four constituent terms: the unique information about the target which is available separately from each explanatory variable alone, the redundant or shared information which is common to the two explanatory variables, and synergistic information, which is information about the target that is available only when both explanatory variables are observed together (e.g. the relationship between their values is informative about the target).^62–64^ In our analysis, the 5 reading scores were used as the target, the features nCTS as the first set of explanatory variables, and behavioral scores as the second set of explanatory variables. PID was also used to better understand the nature of some other statistical associations we uncovered. See Supplementary Methods for further details on PID and its statistical assessment.

## Data availability

The data and the code that support the findings of this study are available at “a link to a OSF repository will be provided upon positive review”.

## 6. Acknowledgements

Florian Destoky and Julie Bertels are (FD) or have been supported by the program Attract of Innoviris (grant 2015-BB2B-10). Julie Bertels is supported by a research grant from the Fonds de Soutien Marguerite-Marie Delacroix (Brussels, Belgium). Xavier De Tiège is Post-doctorate Clinical Master Specialist at the Fonds de la Recherche Scientifique (F.R.S.-FNRS, Brussels, Belgium). Robin A. A. Ince was supported by the Wellcome Trust (grant 214120/Z/18/Z). Mathieu Bourguignon has been supported by the program Attract of Innoviris (grant 2015-BB2B-10), by the Spanish Ministry of Economy and Competitiveness (grant PSI2016-77175-P), and by the Marie Skłodowska-Curie Action of the European Commission (grant 743562).

This study and the MEG project at the CUB Hôpital Erasme are financially supported by the Fonds Erasme (Research Convention “Les Voies du Savoir”, Brussels, Belgium).

## Supplementary Material

### Supplementary methods

#### Recording of video stimuli

The 12 video stimuli of actors telling a story were recorded with a digital camera (Sony handycam, HDR-CX115E). Audio signals were recorded with both the internal microphone of the camera, and an independent high quality microphone (Sony linear PCM recorder, PCM-D50). Audio tracks were synchronized, and only the high-quality audio was kept. Video recordings were framed as a head-shots, and recorded at 50 frames per second (videos were 1920 × 1080 pixels in size, 24 bits/pixel, with an auditory sampling rate of 44100 Hz). The camera was placed ~1 m away from the actors, and the face spanned about half of the vertical field of view. Final images were resized to a resolution of 1152 × 864 pixels. A black old-style-TV-monitor frame was then added to the image.

#### Building the energetic non-informational noise

The (maximally-)energetic non-informational noise was derived from the actual actors’ audio recording by i) Fourier transforming the sound in 2-s-long windows sliding by step of 0.5 s, ii) replacing the phase by random numbers, iii) inverse Fourier transforming the Fourier coefficients in each window, iv) multiplying these phase-shuffled sound segments by a sine window (i.e., half a sine cycle with 0 at edges, and 1 in the middle), and v) summing the contribution of each overlapping window. As a result, the spectral properties of this noise dynamically changed on a ~0.5-s time scale to mirror those of the actor’s voice ~1 s around.

#### Accuracy of speech envelope reconstruction

For each condition and participant, a global value of cortical tracking of the attended speech was evaluated for all left-hemisphere sensors at once, and for all right-hemisphere sensors at once. The decoder tested on a given condition was built based on MEG data from all the other conditions. This procedure was preferred over a more conventional cross-validation approach in which the decoder is trained and tested on separate chunks of data from the same condition because of the paucity of data (*i.e.*, at most ~2.4 min of data per condition). It is based on the rationale that the different conditions do modulate response amplitude but not its topography and temporal dynamics. In practice, electrophysiological data were band-pass filtered at 0.2–1.5 Hz (phrasal rate) or 2–8 Hz (syllabic rate), resampled to 10 Hz (phrasal) or 40 Hz (syllabic) and standardized. The decoder was build based on MEG data from –500 ms to 1000 ms (phrasal) or from 0 ms to 250 ms (syllabic) with respect to speech temporal envelope. Filtering and delay ranges were as in previous studies for phrasal,^28,59^ and syllabic CTS.^47,51,60,61^ Regularization was applied to limit the norm of the derivative of the reconstructed speech temporal envelope,^58^ by estimating the decoder for a fixed set of ridge values (λ = 2^−10^, 2^−8^, 2^−6^, 2^−4^, 2^−2^, 2^0^). The regularization parameter was determined with a classical 10-fold cross-validation approach: the data is split into 10 segments of equal length, the decoder is estimated for 9 segments and tested on the remaining segment, and this procedure is repeated 10 times until all segments have served as test segment. The ridge value yielding the maximum mean RA is then retained. The ensuing decoder was then used to reconstruct speech temporal envelope in the left-out condition. RA was then estimated in 10 disjoint consecutive segments. We then retained the mean of this RA, leaving us with one value for all combinations of subjects, conditions, hemispheres, and frequencies of interest.

Significance of RA in each participant, condition, hemisphere and frequency range was assessed with a *t*-test on the RA values evaluated on 10 disjoint segments.

#### Preprocessing of brain and behavioral indices

All behavioral and nCTS measures were corrected for IQ, age, and time spent at elementary school, and for outliers. For simplicity, we refer to the standardized IQ, age, and time spent at elementary school as the regressors.

An iterative procedure was used to simultaneously control for regressors and deal with outliers. First regressors were regressed out of each measure with an amount of regularization equal to 0.1% the maximal eigenvalue of the regressors’ covariance matrix. Then measures deviating by more than 3 standard deviations from the mean were removed from the distribution. This procedure was repeated until there were no more outliers. Discarded data points were then set to the mean plus or minus 3 standard deviations.

#### Extraction of the relevant features of nCTS

In total, we derived 8 features of nCTS in SiN conditions based on the significant effects of hemisphere and conditions highlighted in Table 3 and Figure 2. There were 4 features for phrasal nCTS: (i) the mean nCTS (the mean standardized nCTS across conditions), (ii) the informational modulation in nCTS (the difference in standardized *nCTS* between informational and non-informational noise conditions averaged across all other factors), (iii) the visual modulation in *nCTS* (the difference in standardized nCTS between *lips* and *pics* visual conditions averaged across all informational noise conditions), and (iv) the hemispheric difference in nCTS (the contrast in standardized nCTS between *left* and *right* hemispheres averaged across all informational noise conditions). The same 4 features were used for syllabic nCTS except that visual and hemispheric modulations were evaluated based on averages across all other factors (and not just across informational noise conditions since visual and hemispheric modulations were seen in all noise conditions for syllabic CTS).

#### Partial information decomposition

PID was previously used to decompose the information brought by acoustic and visual speech signals about brain oscillatory activity,^73^ and to compare auditory encoding models of MEG during speech processing.^62^ As in these references, we measure redundancy with pointwise common change in surprisal for Gaussian variables.^63^ Continuous data values were first subject to a rank-normalisation (copula-normalised)^64^ before being treated as Gaussian variables. A crucial advantage of this redundancy measure as opposed to other PID implementations is that it measures the overlapping information content at the pointwise level and therefore can be interpreted as a within sample (here participant) measure of redundant prediction, directly linked to the coding interpretations of MI. An advantage of the PID over variance partitioning approaches is that unique variance explained might, like conditional mutual information, be confounded by synergistic effects,^123^ whereas PID with common change in surprisal gives the true unique contribution. While the PID has so far been applied within subject to trial data, information theoretic quantities can also be applied as a second level analysis, where each participant is a sample.^124^

The statistical significance of the different information values was assessed with a nonparametric permutation test.^125^ A permutation distribution was computed for each information value by randomly shuffling (10,000 times) children’s target values, and a significance level was computed as the proportion values from the permutation distribution exceeding the observed value.

### Supplementary results

#### Side measures are redundant with RAN and digit span but not with modulations in phrasal nCTS

The information about reading brought by the 3 “side” measures (visual modulation in syllabic nCTS, phoneme suppression and phoneme fusion) was redundant with that brought by a subset (possibly all) of the 4 “main” measures (RAN, phonological memory, visual and informational modulation in phrasal nCTS). To identify this subset, we relied on the PID framework. In this analysis, PID assessed the nature of the information about reading abilities (target) brought by the 3 side measures (first set of explanatory variables) and the 4 main measures (second set of explanatory variables). Specifically, we considered as second set of explanatory variables all possible combinations of the 4 main measures (with or without RAN, with or without forward digit span, with or without the informational modulation in phrasal nCTS, and with or without the visual modulation in phrasal nCTS). Also, the analysis was run separately for the modulation in syllabic nCTS, and for both measures of phonological awareness (phoneme suppression and phoneme fusion) at once. Overall, the results tended to show that the visual modulation in syllabic nCTS provided unique information about reading only with regard to the two phrasal nCTS modulations (see Fig. S1A), and synergic information mainly with the two classical behavioral predictors of reading (see Fig. S1B). Noticeably, the level of synergic information about reading brought by the visual modulation in syllabic nCTS and either or both classical behavioral predictors of reading was not influenced by the addition of either or both of the two phrasal nCTS modulations. Similar results were obtained for the two measures of phonological awareness (see Fig. S1C & D).

### Supplementary discussion

#### Limitations

We did not manipulate the SNR in SiN conditions in the present study. Instead, it was set to 3 dB so that the attended speech was always louder than the noise. Still, one could expect that large effect sizes would be uncovered in more challenging/discriminating listening conditions, as often encountered in classrooms.^126^ However, children’s SiN perception abilities are lower than adults’. Indeed, SiN perception abilities develop until late childhood (≥ 10 years) due to maturation of the auditory system and attentional abilities.^127,128^ Moreover, our data showed that fewer subjects had significant CTS in informational than non-informational SiN conditions for phrasal CTS, and in all SiN conditions for syllabic CTS, with values in the most challenging SiN condition (energetic and informational noise) that were ~50% lower than those in the noiseless condition. Accordingly, setting speech SNR to 3 dB appeared to have made the task challenging enough for children, while ensuring that they could keep their attention focused throughout the experiment.

The amount of data per condition was limited to 2.5 min. Although it may seem little, we evidenced in a previous study that on average, ~30 s of MEG suffice to uncover significant CTS.^28^ Moreover, CTS was significant, when assessed non-parametrically within-participant, in most of our participants in the least challenging conditions (phrasal, 100 %; syllabic, 94%). Of note, more data was used to estimate the regression model mapping MEG data onto reconstructed speech temporal envelope. Indeed, the model used to estimate CTS in each condition was trained on the ~20 min of data from all other conditions. This procedure substantially improved the estimation of CTS compared with a procedure wherein the model was trained and tested in a cross-validation scheme on the data from each condition separately (data not shown). Still, longer data acquisition could have produced more stable CTS estimates, and perhaps stronger associations with reading scores.

## Supplementary tables and figures

**Table S1.**
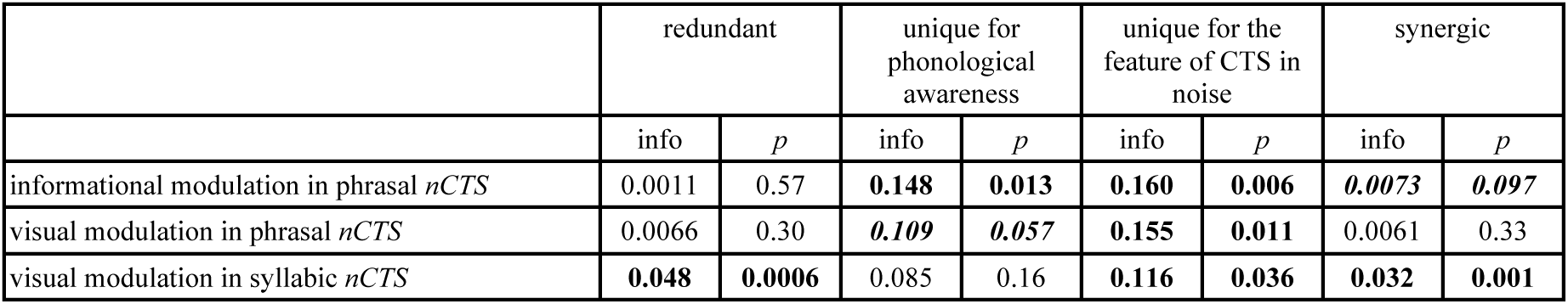
Nature of the information about reading abilities brought by each of the 3 uncovered features of the cortical tracking of speech (CTS) in noise and phonological awareness (mean of the scores for phoneme fusion and suppression). Significant values (*p* < 0.05) are displayed in boldface and marginally significant values are displayed in boldface and italicized.

|  | redundant |  | unique for phonological awareness |  | unique for the feature of CTS in noise |  | synergic |  |
| --- | --- | --- | --- | --- | --- | --- | --- | --- |
|  | info | *p* | info | *p* | info | *p* | info | *p* |
| informational modulation in phrasal *nCTS* | 0.0011 | 0.57 | **0.148** | **0.013** | **0.160** | **0.006** | **0.0073** | **0.097** |
| visual modulation in phrasal *nCTS* | 0.0066 | 0.30 | **0.109** | **0.057** | **0.155** | **0.011** | 0.0061 | 0.33 |
| visual modulation in syllabic *nCTS* | **0.048** | **0.0006** | 0.085 | 0.16 | **0.116** | **0.036** | **0.032** | **0.001** |

**Figure S1.**
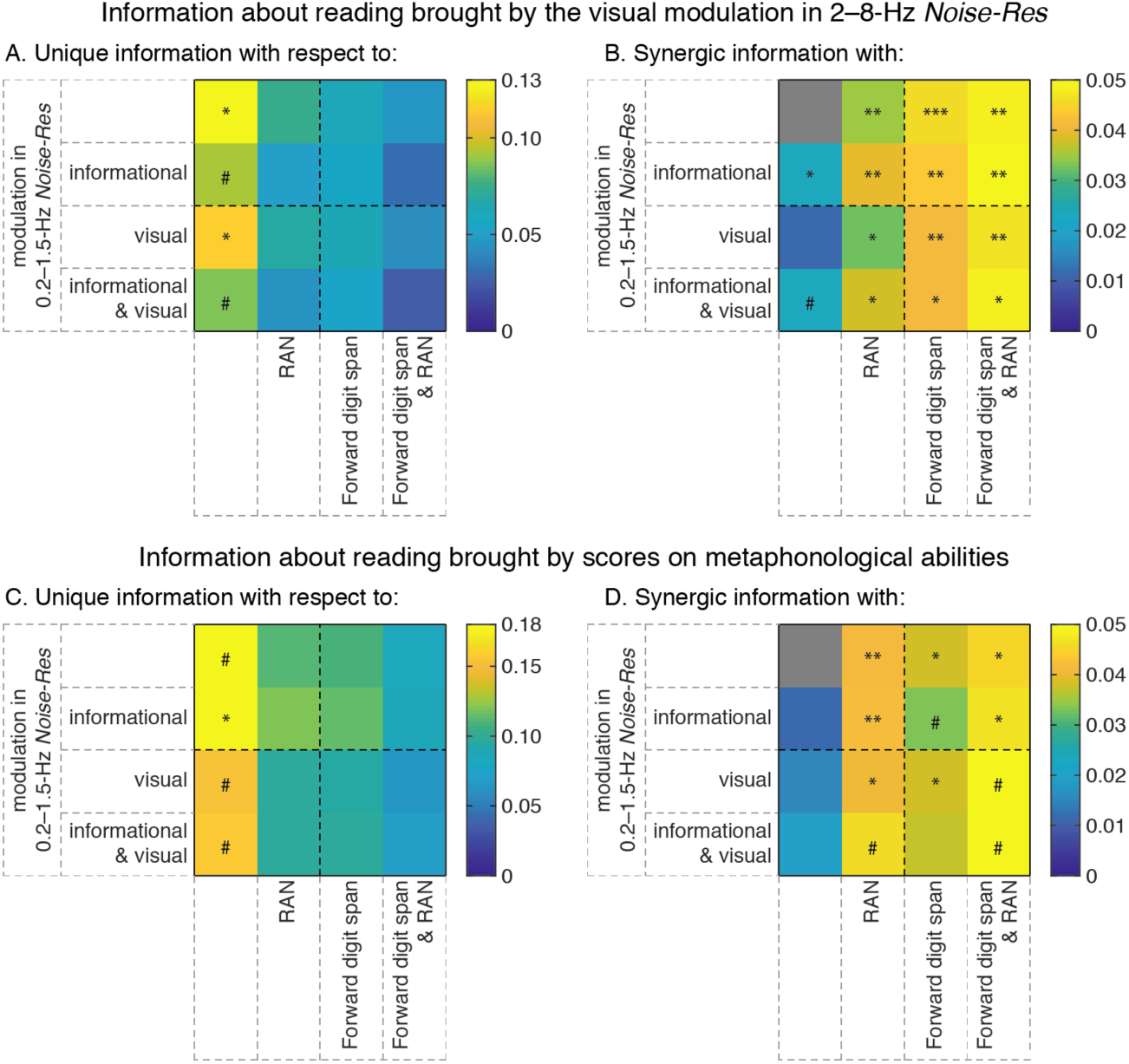
Nature of the information about reading brought by the visual modulation in normalized cortical tracking of speech (nCTS) at syllabic rate (A & B) and metaphonological abilities (C & D). **A** & **C** — Unique information with regard to each possible combination of the 4 regressors included in the final model of reading abilities (without RAN: columns 1 and 3; with RAN: columns 2 and 4; without forward digit span: columns 1 and 2; with forward digit span: columns 3 and 4; without the informational modulation in phrasal *nCTS*: rows 1 and 3; …). **B** & **D** — Same as A for the synergistic information with each possible combination of the 4 regressors. ** *p* < 0.01, * *p* < 0.05, ^#^ *p* < 0.1.

